# HIV Pharmacology Data Repository (PDR): Setting the New Information Sharing Standard for Pharmacokinetic Studies

**DOI:** 10.1101/2023.02.13.528350

**Authors:** Lauren A.R. Tompkins, Oleg Kapeljushnik, Robert Hubal, Adrian Khoei, Julie Dumond, Angela D.M. Kashuba, Alexander Tropsha, Mackenzie L. Cottrell

**Affiliations:** University of North Carolina Eshelman School of Pharmacy, Chapel Hill, North Carolina, USA; Renaissance Computing Institute at UNC (RENCI), Chapel Hill, North Carolina, USA

## Abstract

Clinical therapeutics are becoming increasingly reliant on data science and computational data analytical tools to predict drug behavior in unobserved conditions or populations. These *in silico* approaches, collectively termed “pharmacometrics”, derive biological meaning from the analysis of pooled drug concentration vs. time (CvT) datasets. However, the field lacks standardization for pharmacokinetic (PK) data description and sharing, instead requiring deep dives into the literature and expert data annotation for aggregate data mining. Here we introduce a minimum information sharing standard for PK studies composed of three categories (Intervention, System, and Effects) and implement this standard in the development of a web-based PK database: the HIV Pharmacology Data Repository (PDR). We demonstrate the utility of the HIV PDR by computational modeling of aggregated and curated CvT data extracted from the HIV PDR.

## Introduction

Innovations in computational modeling over the last decade have challenged the old drug development paradigm where pharmacology knowledge is derived from siloed studies with limited applicability outside the context of individual study design. In silico research approaches such as pharmacometrics employ data collected from a wide range of study designs and populations into robust datasets enabling population-level and cross-species inference, as well as systems biology modeling approaches.^123456^ Yet, the rate-limiting step of such innovations is *the ability* to aggregate and harmonize high quality pharmacokinetic data collected over a range of species or populations and anatomical compartments.

Pharmacokinetics is a field of study that follows the path of drugs in the body over time, a process characterized by the acronym “ADME”, referring to drug: Absorption into the bloodstream, Distribution to effector sites such as tissues or distinct cell types, Metabolism (or breakdown) into pharmacologically active and/or inert compounds (metabolites), and Excretion from the body through the kidneys (urine) and/or gastrointestinal tract (feces). The pharmacokinetics of a drug are shaped by characteristics of the drug itself and the system to which it’s administered.^7^ Drug concentrations quantified by the gold standard methodology (liquid chromatography tandem mass spectrometry; LC-MS/MS) over time and in different regions of the body demonstrate the fate of a drug following administration in a pharmacokinetic study. These concentration vs. time (CvT) data, and the basic parameters of studies designed to capture them, naturally lend to acute need to develop and integrated protocols for data collection, harmonization, curation, and integration in a specialized database that could facilitate data sharing and modeling.

The trend toward data collection and consolidation into specialized databases is an undeniable attribute of modern scientific research in nearly any biomedical discipline as evidenced by rapid proliferation of the principles of data science across basic and applied research.^8^ Multiple databases have been created over the years and many critical databases that support an accelerate biomedical research have been supported by the National Institutes of Health.^9^ In recent policy documents, this major funding agency mandated that biomedical researchers should “promote the management and sharing of scientific data generated from NIH-funded or conducted research”. To support data sharing, NIH created a list of NIH supported public sharing resources ^10^ that investigators can use to share their data. The proposed list is extensive but, naturally, not exhaustive, so there is an expressed need to establish novel public data repositories that can facilitate data sharing by scientists working is specialized disciplines where NIH supported depositories may not be most adequate. Arguably, pharmacokinetics is an example of such discipline as the search of the aforementioned NIH resources^10^ with the key word “pharmacokinetics” produced no hits.

Here we introduce a minimum information standard for PK studies generating CvT data in humans and animals. The concept of reporting *minimal information standards* has been gaining popularity in different research disciplines.^11^ Notable examples include Minimum information about a bioactive entity (MIABE)^12^, Minimum information about a microarray experiment (MIAME)^13^, or Minimal Information About Nanomaterials (MIAN).^14^ The new standard we propose here consists of key reportable components divided into 3 categories following the operative structure of PK studies: intervention (drugs administered, including how and when they are administered at a given quantity), system (species the intervention is administered to, including anatomical compartments being sampled), and effects (small-molecule concentrations measured upon intervention, including prodrugs, drug metabolites, and endogenous markers). We then apply this standard in curating PK data collected from previously siloed studies into a user-friendly HIV PDR database to support data sharing, management, and mining across the pharmacokinetics and pharmacometrics community. We illustrate the power of PDR by a proof-of-concept computational modeling of CvT data facilitated by the availability and accessibility of respective data in PDR.

### Categories of the Minimum Information Standard for PK Studies

#### Intervention

The Intervention category sets the stage for a PK study, capturing the drug being administered, the quantity given and how (i.e., dose and route of administration; Table 1). These key reportable components, the “pharmaco” portion of a PK study, guide interpretation of resulting drug concentrations and thereby feed into ADME characterization from absorption all the way to excretion. Additionally, data surrounding time, the “kinetic” portion, are equally critical for determining the onset, duration, and intensity of a drug’s effect. Thus, the Intervention category also includes three components related to time: time elapsed since administration of a dose (for generating, or interpreting data via, a PK curve), frequency of administration and medication start date (for identifying the probability that a steady-state condition has been achieved), and formulation attributes to delineate between immediate- or extended-absorption profiles.

**Table 1:** Minimum Information Standard for PK Studies.

| **Category and Component** | **Units** | **Requirements and Rationale** |
| --- | --- | --- |
| ***Intervention*** |  |  |
| Administered Drug | n/a | Generic name, with brand name provided for coformulated pills |
| Administered Dose | mg, mg/kg, ug/kg | Human/animal weight required when mg/kg or ug/kg used |
| Route of Administration | n/a | Full route spelling rather than abbreviations to prevent ambiguity |
| Frequency of Administration | Hours (qXXh), Once, Baseline, Placebo/Sham | qXXh nomenclature rather than b.i.d., etc. to prevent ambiguity |
| Weight (human/animal) | kg | Inclusion for PK computation and dose calculations (where needed) |
| Time Elapsed Since Last Dose | Days, Hours, Minutes | Consistent use across all PK study samples |
| Medication Start Date | n/a | MM/DD/YYYY format to prevent ambiguity |
| ***System*** |  |  |
| Species | n/a | Controlled vocabulary and hierarchies in Figure 1 |
| Matrix | n/a | Controlled vocabulary and hierarchies in Figure 1 |
| ***Effects*** |  |  |
| Analyte | n/a | Matched to nomenclature used in Administered Drug |
| Analyte Concentration | ng/mL, ng/g, g, fmol/million cells | Values included for any conversion calculations to yield units employed |

For PK studies in plasma, the Intervention category reports essential data required to generate a drug CvT profile (or PK curve) to calculate the basic PK parameters of a drug: Cmax (maximum observed concentration), Tmax (time it takes to reach Cmax), AUC (area under the curve reflecting total exposure to the drug), and half-life (time it takes for half the drug to be eliminated). These parameters indicate how quickly the drug enters the peripheral blood (absorption, “A” in ADME) and leaves it for distribution, metabolism, and excretion (“DME” in ADME).^7^

#### System

The System category contextualizes a PK study within the physiological properties of the species employed, including the anatomical compartments sampled by collection of distinct biological matrices (fluids, tissues, cell types; Table 1 and Figure 1). Preclinical studies in animals permit sample collection of a variety of matrices obtained at discrete time points following administration of a drug in a controlled environment. Likewise, clinical or exploratory studies in humans define the basic PK parameters associated with a protective effect in end-user populations and may involve measurement of drug concentrations in different regions of the body, often at effector sites where the drug enacts an intended therapeutic objective (indication).^15^ Data obtained in either type of PK study can be queried to inform the other through *in silico* modeling such as for scaling up studies from small animals to nonhuman primates to humans in drug development programs^9^ or for repurposing existing FDA-approved drugs for alternative therapeutic uses investigated in animal models. Examples of the latter include adapting oncology agents to reverse latent infection in HIV cure approaches^16,17^ or applying ritonavir, a PK boosting agent developed for HIV, to treat SARS-CoV-2 when administered with nirmatrelvir (ie, PaxlovidTM; Pfizer).^18^

**Figure 1.**
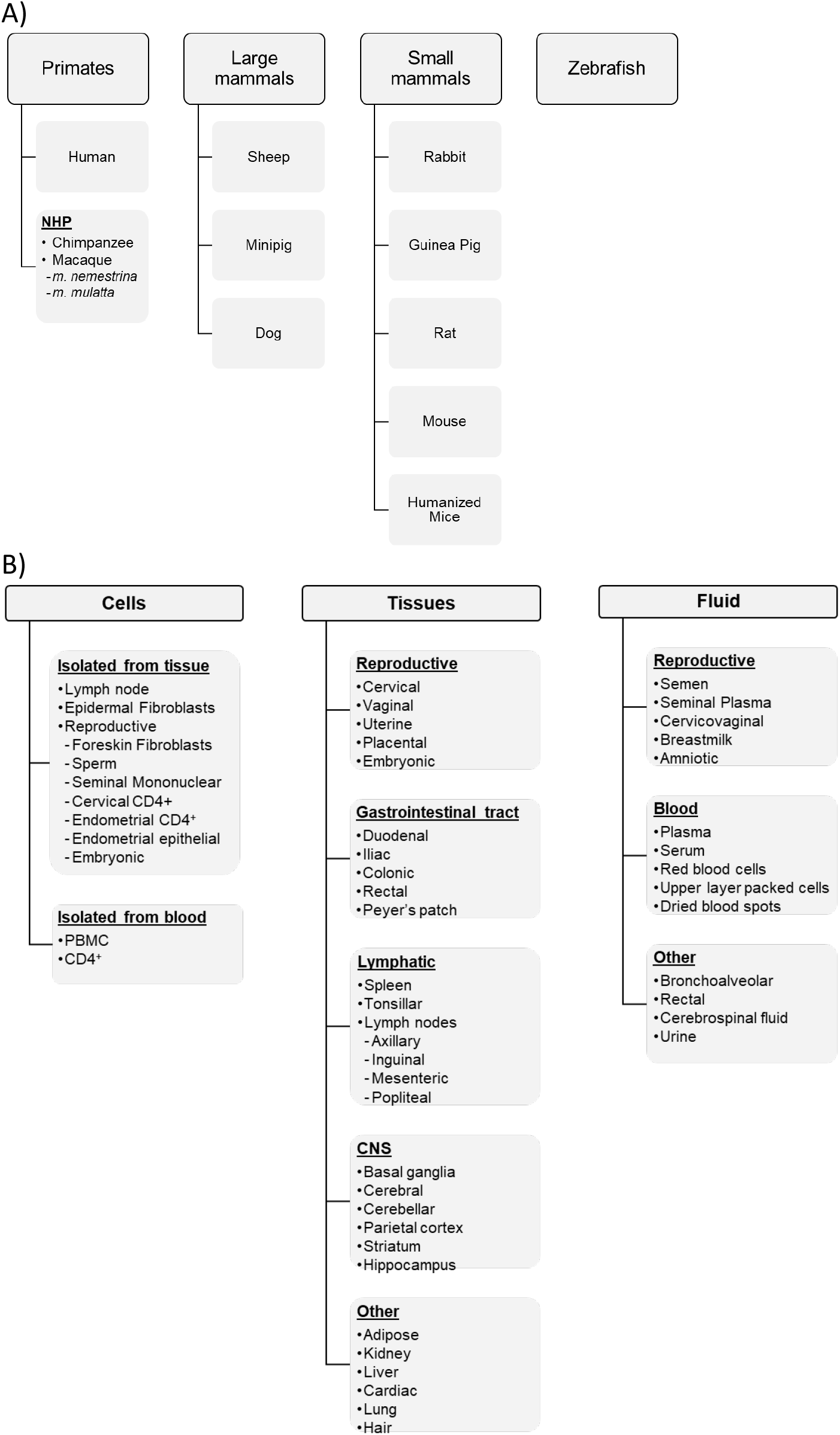
Taxonomical and anatomical hierarchies proposed for the minimum information standard for PK studies. The species (A) and anatomical source (B) associated with CvT data are delineated into field-relevant hierarchies. Additional variables may be incorporated into the basic structure suggested.

Direct quantification of drugs and drug metabolites in distinct biological matrices illustrates drug distribution to sites of effector function and metabolism, including sites where prodrugs are converted to pharmacologically active moieties. Thus, individual PK studies are often designed to characterize drug distribution in specific anatomical compartments (eg, CNS, GI, genital) to answer specific questions about therapeutic effect or toxicity. Physiologically based PK modeling draws inferences about how these compartments relate as drugs flow through the body.^19,20^ By defining hierarchies of biological matrices that map on a whole-body scale across species (Figure 1), the System category relates drug concentration data to meaningful keywords for data queries that support these types of modeling activities.

#### Effects

The goal of defining a minimum information standard for PK studies is to parse drug concentration data for which these studies are conducted. The Effects category coalesces concentrations of prodrugs, drug metabolites, and other small molecules such as biomarkers into two standardized components: Analyte and Analyte Concentration (Table 1). Linking these concentrations with variables under the Intervention and System categories generates a minimum information standard with the capacity to support biologically meaningful data queries. The standard incorporates data for mechanistically related molecules to allow comparisons of a drug’s metabolic pathway, such as from prodrug to drug metabolite to active moiety, with its effector function, such as target substrates for competitive inhibition. The flow of drugs can be followed in time from absorption all the way to urinary or gastrointestinal excretion in different species and following distinct routes of administration.

### Piloting the New Standard with the HIV Pharmacology Data Repository

#### HIV Pharmacology as a Model System for PK Database Generation

One approach to ensuring a minimum information standard balances minimalism with broad applicability is to trial the standard with a complex system. Antiretroviral therapy (ART) involves complex pharmacological issues identified and addressed by over three decades of HIV treatment evolution.^21,22,23^ Studies in HIV pharmacology have pioneered or advanced concepts such as U=U (Undetectable=Untransmittable by sexual intercourse), also known as treatment as prevention (TasP),^24,25^ pre-exposure prophylaxis (PrEP),^26,27^ and pharmacokinetic enhancement with boosting agents.^28^ Yet, despite broad implementation of effective therapy capable of sustaining virologic suppression in people living with HIV and preventing infection in high-risk populations, annual incidence continues to drive the global burden of disease.^29^ The success of pharmacological strategies to reduce HIV incidence relies on high adherence to daily oral dosing in people living with, and at risk for, HIV who may not be well-positioned for success due to lack of access to care, low health literacy, and/or discrimination and stigma.^303131^ Efforts to reduce pill burden through the advent of long-acting formulations are only beginning to come to fruition with FDA approval of the first such complete ART regimen (ie, injectable Cabenuva, cabotegravir plus rilpivirine; ViiV Healthcare).^32,33^ Additionally, strategies to cure HIV are complicated in part by pharmacological challenges (eg, limited drug distribution at effector sites such as in tissues, inadequate therapeutic efficacy, and unfavorable drug toxicity).^34,35^

#### Quality of Bioanalytical Data

The clinical pharmacology field has recognized the need for standardization in bioanalytical quality control to ensure scientific rigor and reproducibility across laboratories generating drug concentration data for clinical studies. To this end, the AIDS Clinical Trials Group (ACTG) established a comprehensive quality assurance/quality control program, including routine proficiency testing consistent with Clinical Laboratory Improvement Amendments of 1988 (CLIA) requirements for clinical tests.^36^ The Clinical Pharmacology Quality Assurance (CPQA) program was established in 2008 to expand on these efforts by enforcing bioanalytical standardization, including passing criteria for instrument runs, across participating laboratories (currently 12) to ensure LC-MS/MS methods consistently yield accurate data values over time.^37,38^ Yet, the pharmacological context from which those values were derived is not documented, representing a missed opportunity for the CPQA network. Implementation of the minimum information standard for PK studies proposed herein would harmonize large drug concentration datasets with meaningful biological context required for aggregate PK modeling exercises.

#### The World’s First HIV Pharmacology Data Repository

As a CPQA-participating laboratory and Core facility of the UNC Center for AIDS Research (CFAR), our laboratory is uniquely positioned to trial the minimum information standard for PK studies proposed herein using diverse preclinical and clinical CvT datasets generated from over 20 years of bioanalytical CFAR service provision. To this end, we have consolidated our electronic data archives into a central, searchable HIV Pharmacology Data Repository (HIV PDR). This interoperable data portal uses a relational database structure (on an SQL platform) to integrate the minimum information standard for PK studies with drug concentration data generated according to field-established quality standards. Importantly, the HIV PDR was designed according to FAIR principles for data sharing^39^ and curated according to HIV pharmacology nomenclature to bypass the need for literature deep dives and ultimately drive user-driven *in silico* research.

Currently, about 62% of electronically archived data generated from CFAR service requests are maintained in a database stored in an SQL server with the front-end developed using an ASP.NET core with Angular and the back-end on an SQL Server 2017. Our initial developmental activities collapsed 924 unique column headers describing bioanalytical results within 610 Excel files into 145 unique bioanalytical variables, thereby populating the *Effects* category of information described above. Additional data dictionaries were created to harmonize terminology for sample attributes (ie, *Intervention* and *System* categories) that are transmitted by CFAR service requesters through the manifest Excel file. In creating these dictionaries, we collapsed 246 descriptors of sample attributes into 80 variables of unique anatomical sources obtained from 15 species, with taxonomical and anatomical hierarchies employed (Figure 1). The result of these development efforts is a searchable database with controlled vocabularies housing >75,000 unique drug CvT values resulting from >29,000 samples collected across investigations of 72 drug molecules (Figure 2). Subsequent harmonization efforts will liberate an additional >40,000 CvT datapoints archived in approximately 350 electronically archived files generated prior to 2016 and requiring minor reformatting for upload.

**Figure 2.**
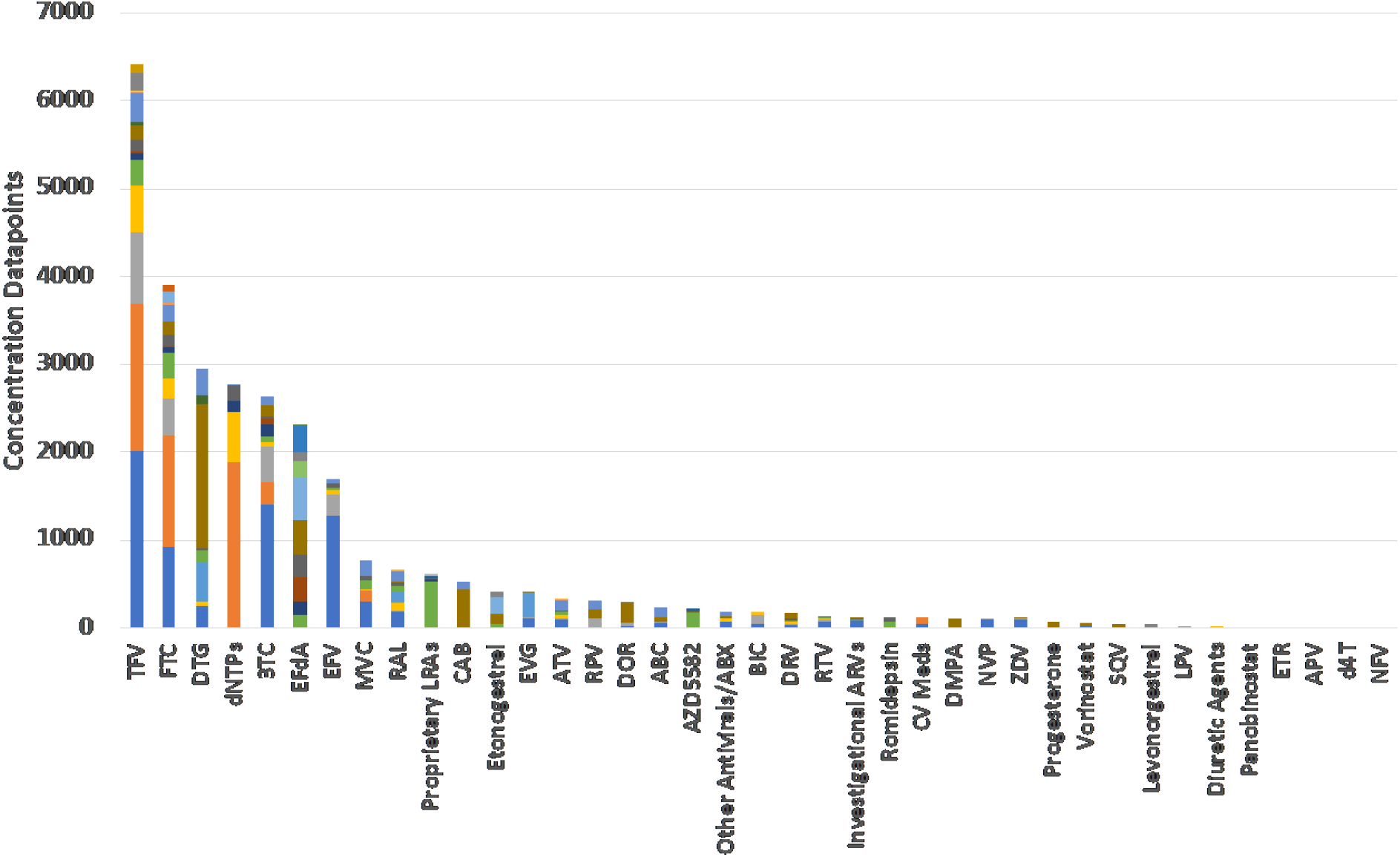
Summary of aggregate CvT values injected into the HIV PDR as of January 2023. Bar colors indicate unique species and matrices from which values were derived.

To minimize unnecessary expansion of HIV PDR dictionaries, we re-envisioned our template manifest Excel file to capture relevant information based on 4 distinct categories of pharmacology studies: clinical PK, preclinical PK, adherence monitoring, and in vitro/ex vivo studies. These new file structures harness cell validations that constrict entries for sample information using categorical values already contained within our dictionary and apply meaningful formats for numerical values such as dose, time, date, or animal weight. Thus, a controlled vocabulary is enforced with constrained options to reduce the need for data curation, minimize term ambiguity, and improve searchability for data mining and artificial intelligence (AI) in the future. These cell validations can be easily updated with new variables if expansion of our existing dictionary is needed for a new service request.

#### Extractable Data Queries Promote Scientific Exploration

Datasets are extractable from the HIV PDR as machine-readable CSV files that can be tailored for a particular study question through employing a series of filters including species, biological matrix, and drug. Hierarchical classifications can be employed for targeted searches on an anatomical level. Hierarchies distinguish bodily compartments and the subcompartments within to aggregate data obtained from an entire organ (eg, GI tract) or individual areas (eg, Peyer’s patches, upper GI, lower GI), with delineation at the cellular level (eg, vaginal CD4+ cells vs. vaginal epithelial cells) and fluid level (eg, peripheral vs. anatomical site).

As a proof-of-concept demonstration of scientific utility, we performed a secondary data analysis to develop a population PK model for the prodrug tenofovir disoproxil fumarate (TDF) by extracting 3941 CvT values for tenofovir (TFV, the drug metabolite in circulation) resulting from human plasma samples. Then we applied additional filters to exclude TFV CvT data points associated with an alternative prodrug, tenofovir alafenamide (TAF). These filters resulted in a CvT dataset containing 947 observations from 100 human study participants across 3 dosing levels of TDF under first-dose and steady-state conditions (Figure 3). We successfully fit a two-compartment population PK model to this dataset using the PK modeling software NONMEM without any data formatting related errors. Interindividual variability was estimated on the clearance and volume of central compartment parameters, and the model parameters were estimated with reasonable precision (CV% < 8.81). The model structure and parameter values are consistent with existing literature^.40^

**Figure 3.**
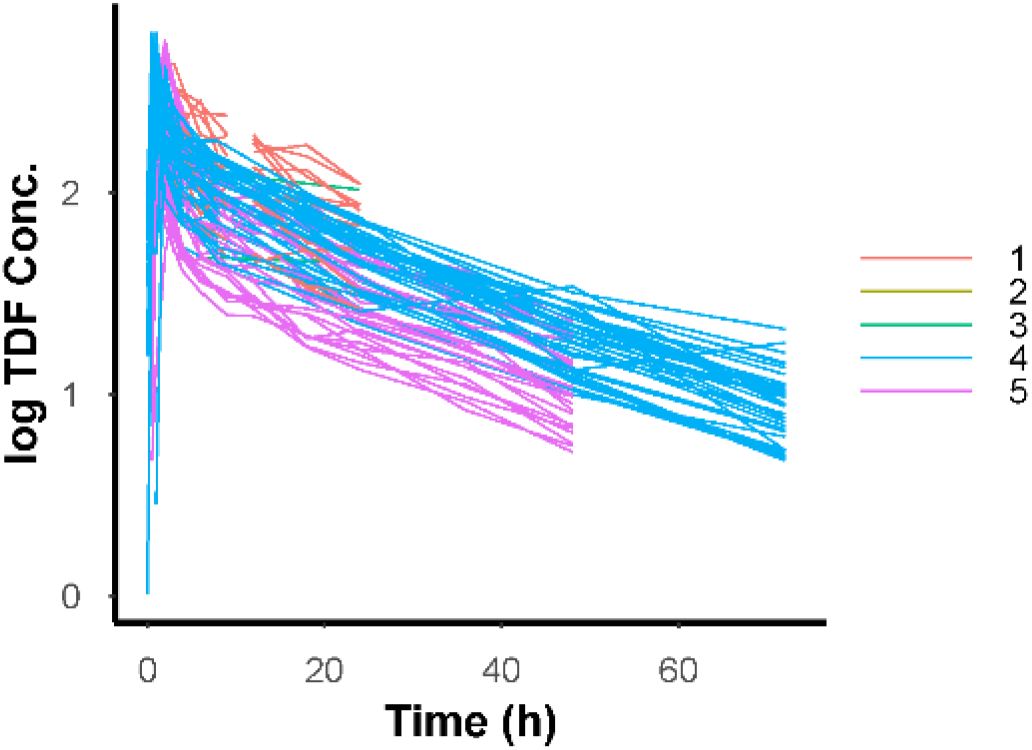
Human PK dataset extracted from the HIV PDR. 100 individual human TFV plasma PK profiles following TDF dosed orally at 150, 300, and 600mg aggregated across 5 studies (study categorized by color; n=947 observations).

## Discussion

Here we propose a minimum information standard for PK studies, using HIV pharmacology as a model system, and apply this standard in the design of a web-based data portal to support *in silico* research driving next-generation therapeutics (the HIV PDR). The proposed standard was intentionally designed to be truly *minimal* following the basic tenets of pharmacokinetics. The basic structure of the standard, which is divided into three categories (Intervention, System, and Effects), allows flexibility to include additional components if supported by demonstrated potential for broad applicability. Applying the minimum information standard through the HIV PDR provides the opportunity to establish suitability and determine areas of enhancement through end user engagement. The use of standard data annotation and ease of access and focused data extraction can drastically facilitated multiple data modeling and benchmarking studies.

The HIV PDR innovates in the field of clinical pharmacology, as existing platforms (Drugs@FDA^41^ and HIVDrugInteraction^42^) are limited in pharmacometric utility. These platforms archive summary estimates of PK/PD parameters housed within the drug label rather than raw extractable CvT data. Furthermore, these repositories lack standardization in parameter reporting and provide data exports in nonmachine readable format, creating bottlenecks in the model development process. Large clinical cohort networks such as MACS/WHIS^43^ and ACTG^44^ may aggregate CvT data archived within the network, but they lack minimum information standards and require lengthy application processes that hinder the accessibility of these data pools.

Importantly, existing clinical pharmacology databases have little opportunity to share data generated in real-time for investigational pharmaceutical agents. Most of the data provided is sourced directly from the new drug application (NDA) or the product label. The HIV PDR overcomes this limitation through synergy with existing mechanisms for PK data stewardship within the CFAR network. As a generator of bioanalytical data that participates at every phase of the drug development process, our CFAR Core facility has established processes for streamlining information transfer for investigational compounds assessed in federally funded studies. For example, the HIV PDR houses >3,500 concentrations of islatravir (a first-in-class, investigational antiretroviral) and its active metabolite in plasma, cells, and tissues from mice, rats, rabbits, dogs, sheep, and nonhuman primates (NHPs) following intravenous (IV), subcutaneous (SQ), and oral administration. Collating and sharing data from these federally funded projects in the HIV PDR offers an unprecedented opportunity to conduct modeling studies that support dose translation and obviate the risk of research duplication.

The 2023 Final NIH Policy for Data Management and Sharing (NOT-OD-21-013) strongly encourages that all scientific data resulting from NIH-funded research be shared through and preserved in established repositories.^45^ This recommendation demonstrates the preference for a consolidated approach to data sharing over an institutionally siloed one; yet, established PK data repositories do not exist for the HIV/AIDS research community. Based on a comprehensive search of the indexing sites, FAIRSharing and PubMed, we are only aware of two existing PK data repositories. One archives 16,267 CvT datapoints for 187 chemical entities and their metabolites.^46^ However, the repository’s rigid upload criteria (requiring that data be reformatted into a template form) and their environmental toxicokinetic focus make this repository an unlikely site of data sharing for HIV researchers. The other database archives 73,017 CvT datapoints for 10 pharmaceutical agents.^47^ However, the database is constrained to clinical plasma PK data unrelated to HIV and is extremely limited in the breadth of archived drugs (acetaminophen, caffeine, codeine, diazepam, glucose, midazolam, morphine, oxazepam, simvastatin, or torasemide).

### Future Directions in HIV PDR Operational Efficiency

The HIV PDR is designed for growth in keeping with research strategies executed through CPAC Core activities and beyond. Future design activities include a web-based CFAR service request system that utilizes an algorithm-based questionnaire to obtain study information that is parsed into an Excel spreadsheet containing data input categories tailored to the specific study described. The resulting *study template* will prompt for relevant components of the minimum information standard for each sample associated with CvT data. To accommodate situations in which completing a study template may be prohibitively time-consuming or otherwise challenging, such as injection of archived data from external bioanalytical laboratories, the HIV PDR can apply existing synonym dictionaries to parse data presented differently for reclassification into the minimum information standard the system. In initial activities developing the HIV PDR, Excel file structure rules were designed to facilitate submission of archived files from a wide range of sources, where only a minimum set of criteria (currently 2 requirements) need to be met for automatic injection. Thus, minimal file reformatting may be anticipated to meet these criteria.

Another high priority opportunity includes designing data validation tools within the HIV PDR that flags drug concentration values outside of the normal distribution for existing data. These tools can be designed to match newly generated datapoints against historical controls according to the following variables: analyte, concentration units, biological matrix, species, drug, dose, and route of administration. The validation tool would build on existing quality control standards to add another level of quality assessment prior to data publication. Finally, an accession system that records data query attributes and harnesses system-versioned temporal tables to ensure each extracted PK data pool will be an important design feature to ensure reproducibility despite the dynamic nature of the data archives. In keeping with the new NIH policy for data sharing, this accession system will provide a persistent unique identifier to support proper citation in the literature and track changes in scientific trends as new CvT data becomes available over time.

## Conclusion

Here we propose a minimum information standard for PK studies. The standard was designed to be truly *minimal* and divided into three essential categories (Intervention, System, and Effects) to derive pharmacokinetic meaning from bioanalytical data. By applying this standard, we successfully incorporate >75,000 CvT datapoints for 72 drug molecules archived in ∼350 electronic files into a searchable database, the HIV PDR. Finally, we demonstrate utility of the HIV PDR to promote scientific exploration by using the database to extract TFV PK data and perform population PK modeling. To our knowledge this is the first report of its kind within the HIV pharmacology field, and we believe our approach to data management represents a high priority opportunity within the field to facilitate user-driven in silico research.

